# Guiding Interoperable Electronic Health Records Through Patient Sharing Networks

**DOI:** 10.1101/473272

**Authors:** Jonathan M Clarke, Leigh R Warren, Sonal Arora, Mauricio Barahona, Ara W Darzi

## Abstract

Effective sharing of clinical information between care providers is a critical component of a safe, efficient health system. National data sharing systems may be costly, politically contentious and do not reflect local patterns of care delivery. This study examines hospital attendances in England from 2013 to 2015 to identify instances of patient sharing between hospitals. Of 19.6 million patients receiving care from 155 hospital care provider, 130 million presentations were identified. On 14.7 million occasions (12%), patients attended a different hospital to the one they attended on their previous interaction. A network of hospitals was constructed based on the frequency of patient sharing between hospitals which was partitioned using the Louvain algorithm into 10 distinct data-sharing communities, improving the continuity of data sharing in such instances from 0% to 65-95%. Locally implemented data-sharing communities of hospitals may achieve effective accessibility of clinical information without a large-scale national interoperable information system.

## Introduction

As modern healthcare systems move towards service centralisation^1–4^, the care of individual patients is increasingly shared between several providers. Healthcare providers involved in patient sharing should coordinate their efforts to provide effective, integrated care across the patient journey. However, providers may sometimes operate as ‘silos’, often without knowledge of the problems addressed, services provided or medications prescribed by the previous hospital^5,6^. The incomplete exchange of health information during transitions of care may lead to ineffective care^7^, adverse patient outcomes and additional healthcare spending^8^. Rather than promoting a seamless care continuum, patient sharing can therefore lead to care fragmentation, particularly for patients with multiple co-morbidities and complex-care needs.

Previous studies have demonstrated that breakdown in communication between healthcare providers is the leading root cause of sentinel events in hospitals^9^. Enhancing communication during transitions of care could therefore improve patient safety and reduce healthcare costs ^10^. Understanding when and where patients access care is an essential step in establishing effective communication strategies. Identifying patient sharing between healthcare organisations may inform the development of effective, efficient data-sharing practices at local, regional and national levels ^7^.

Achieving effective interoperability between health providers has become a priority for health systems across Europe and the United States ^11–13^. Despite the apparent benefits to the safety and efficiency of care provision, interoperability in health systems in the US lags behind expectations ^11^. Attempting to achieve large scale interoperability between care providers nationally across large health systems may be complex, expensive and politically contentious ^11,14,15^. Such strategies also go against common patterns of patient care. Healthcare is a locally delivered phenomenon and where care is shared between hospitals, those hospitals are often spatially proximate. Establishing local or regional strategies for interoperability may therefore provide similar coverage to a nationally interoperable EHR system in a more cost-effective, socially and politically palatable manner.

Interactions between patients and healthcare providers are critical to understanding the behaviours of modern health systems ^16^. Recent decades have seen the application of techniques from the field of network analysis to understand a range of complex, dynamic social systems ^17–19^. In the healthcare context, network analysis approaches may emphasise the connections that exist between healthcare providers and patients. In doing so, they provide insights into the complex interconnections that exist within modern health systems that would only be apparent through an evaluation of the system as a network.

One method of studying patient sharing between providers or organisations is through the creation and analysis of *patient-sharing networks*^20–22^. Such networks provide a representation of the links between healthcare providers, based on patients they share^23–26^. This approach may be extended to identify occasions where patients attend one healthcare provider having attended a different healthcare provider on the last time they received hospital care. These episodes identify occasions where the information available to one clinician may be incomplete or outdated and therefore compromise clinical care ^27^. It is then possible to identify how these patterns of patient sharing and disrupted clinical data flows are distributed geographically and across hospital trusts. Insights from these findings may be used to inform and improve evidence-based healthcare policy.

Our objectives are: (1) To use outpatient, inpatient and accident and emergency admission Hospital Episode Statistics (HES) data, to identify inter-organisational patient-sharing connections in the NHS in England. (2) To use the Louvain modularity community detection process to partition this patient sharing network into discrete, mutually exclusive, collectively exhaustive data-sharing communities to guide hospital interoperability.

## Results

In the year 2013-14, 19,682,360 unique patients were identified attending a total of 155 acute hospital trusts. In the 12 months from their first recorded presentation, these 19.7 million patients interacted with secondary care a total of 130,161,023 times, including their first presentation. 16,002,415 of these patients (81.3%) had more than one recorded interaction with secondary care. Of those presenting more than once, 4,162,780 patients (26.0%) presented to more than one secondary care provider.

126,481,078 (97.2%) secondary care interactions were recorded for patients that presented on more than one occasion over the 12-month period. Of these, 46,966,969 (37.1%) pertained to patients who presented to more than one provider. On 14,748,791 occasions (11.7%), patients who presented more than once in secondary care had immediately previously attended another provider.

Figure 1 maps the probability that a patient resident in each LSOA attends different providers on consecutive interactions during the study period. This measure characterises, at the population level, the probability of patient-sharing across secondary care providers. The probability of presenting to different providers ranges from 0% to 30%, with a higher probability in urban areas with greater spatial density of providers; on the other hand, East Anglia, Cornwall and East Yorkshire have particularly low probabilities of presentation to different providers.

**FIGURE 1.**
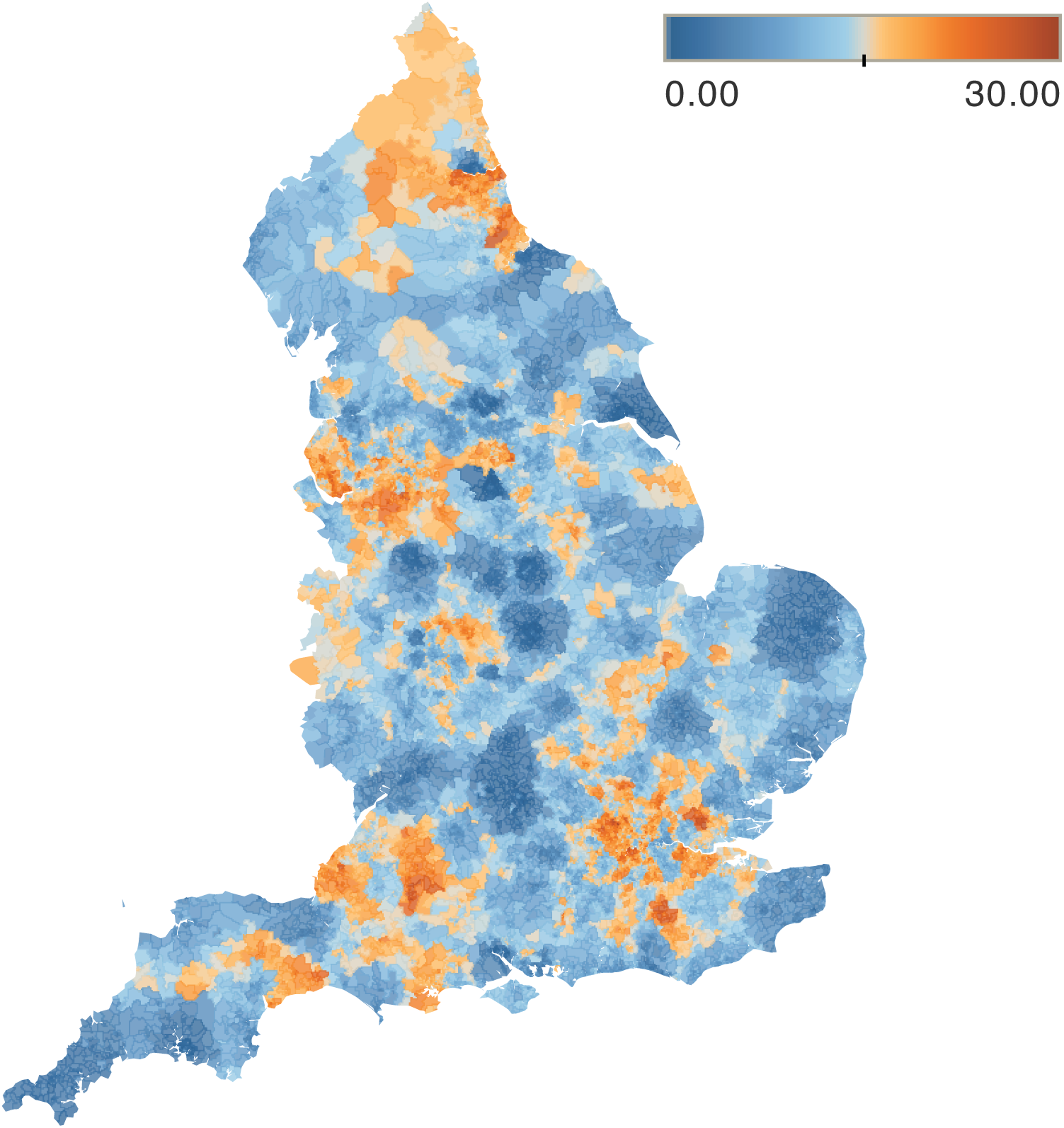
Spatial distribution of patient sharing. Geographic representation by Lower Layer Super Output Area (LSOA) of the probability (as represented by the colour map) that a patient presenting to one provider was last seen at a different provider within the study period.

Patient sharing is extensive: the patient-sharing network between the 155 providers has a total of 11,641 edges present (97.5% out of a possible 11,935 edges) indicating a densely connected network. However, the magnitude of patient sharing is highly inhomogeneous across pairs of providers: only 1034 edges (9.1%) have a weight of more than 1000 cases, suggesting that each hospital mostly shares patients with a small number of other hospitals. Figure 2 shows two representations of the patient-sharing network, made sparser for clarity of illustration, showing edges of weight above 100 cases (left) and edges of weight above 10,000 cases (right).

**FIGURE 2.**
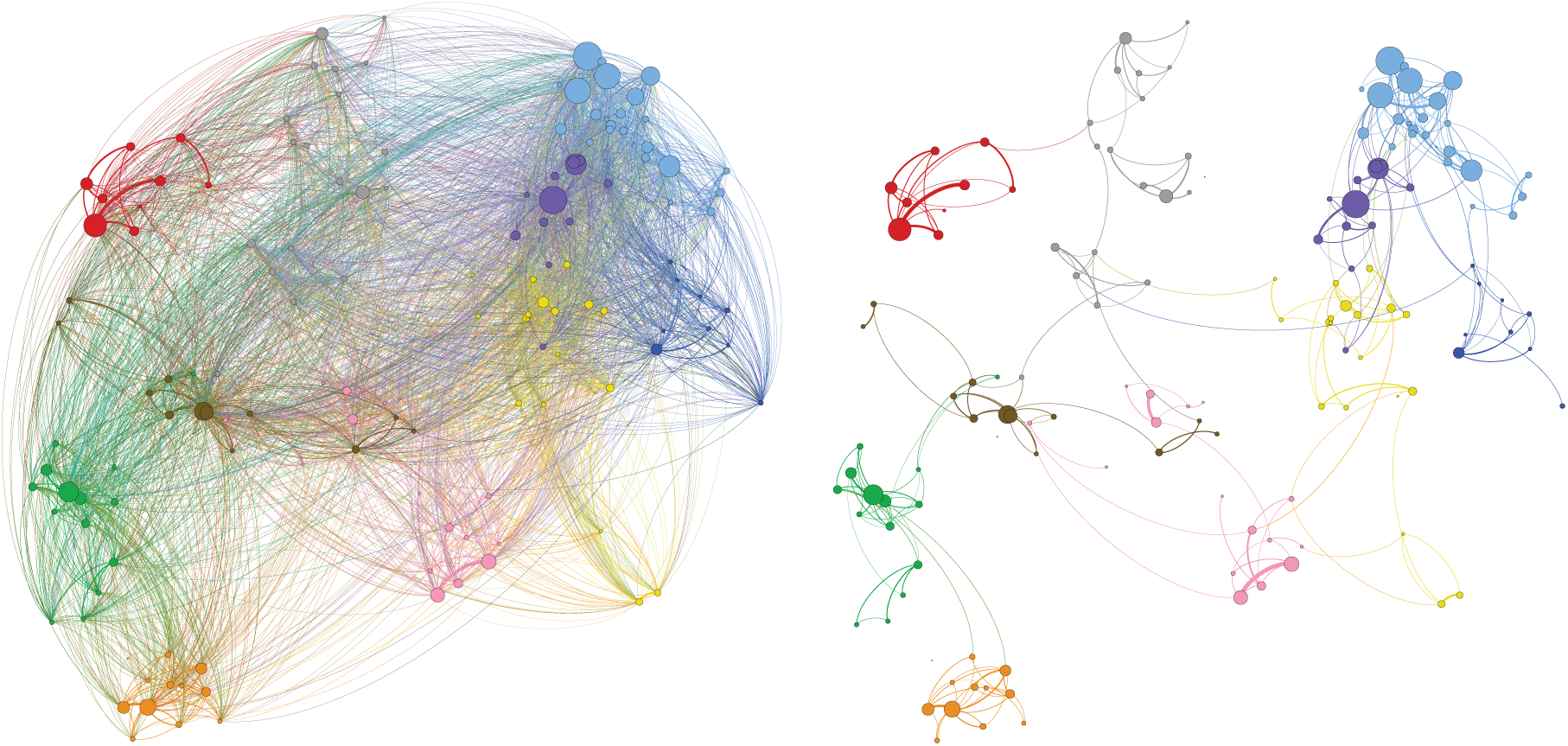
Interhospital patient sharing networks. Two representations of the patient-sharing network of healthcare providers made sparser for illustration purposes. The network has edge weights given by the patient-sharing cases (*F*_*AB*_) and the size of the nodes reflects the total number of shared patients. Left: patient-sharing network with edges above 100 cases shown (4397 edges – 38.6%). Right: patient-sharing network with edges above 10,000 cases (346 edges – 3.0%).

The full patient-sharing network was partitioned into data-sharing communities through Louvain optimisation of resolution modularity (19). The analysis generated 10 stable communities of providers with a modularity value of 0.818, suggesting good partitioning. The 10 communities obtained contain between 9 and 24 care providers, and the frequency of episodes where patients present to one provider having previously attended another provider within and outside the community are shown in Table 1. The nodes (health providers) in Figure 2 are coloured according to their community of membership, and the size of each node is proportional to the sum of transfer flows *F*_*AB*_ for the hospital it represents.

**TABLE 1.** Data sharing community descriptions. Table showing for each interaction with a provider in each community the proportion of times the patient’s previous secondary care interaction had been with a provider located inside or outside the community of that provider.

| Community Name | Number of Hospitals | Total Fragmented Presentations | Location of Previous Provider |  | % Within Community |
| --- | --- | --- | --- | --- | --- |
|  |  |  | Within Community | Outside Community |  |
| *'East Anglia'* | 10 | 537,408 | 385,490 | 151,918 | 71.73 |
| *'Merseyside'* | 12 | 1,002,066 | 888,043 | 114,023 | 88.62 |
| *'North East'* | 9 | 1,086,068 | 1,024,145 | 61,923 | 94.3 |
| *'North London'* | 24 | 3,373,655 | 2,696,511 | 677,144 | 79.93 |
| *'North West'* | 15 | 1,476,174 | 1,283,869 | 192,305 | 86.97 |
| *'South'* | 16 | 1,157,703 | 754,640 | 403,063 | 65.18 |
| *'South London and South East'* | 16 | 2,107,474 | 1,567,297 | 540,177 | 74.37 |
| *'South West'* | 16 | 1,172,622 | 986,142 | 186,480 | 84.1 |
| *'West Midlands'* | 16 | 1,343,370 | 1,137,792 | 205,578 | 84.7 |
| *'Yorkshire and North Midlands'* | 21 | 1,492,251 | 1,256,115 | 236,136 | 84.18 |

Figure 3 represents the geographical location of each provider coloured according to their community of membership. This figure shows that the communities found from the patient-sharing network in an unsupervised manner have a clear geographical and regional pattern, which, crucially is not imposed *a priori*, but rather results directly from the data reflecting patterns of patient sharing that favour proximity between providers. In turn, the analysis reveals regions where patients are more likely to be shared within a defined group of providers, rather than outside of it. This phenomenon is again a consequence of patterns of patient behaviour rather than *a priori* defined administrative groupings.

**FIGURE 3.**
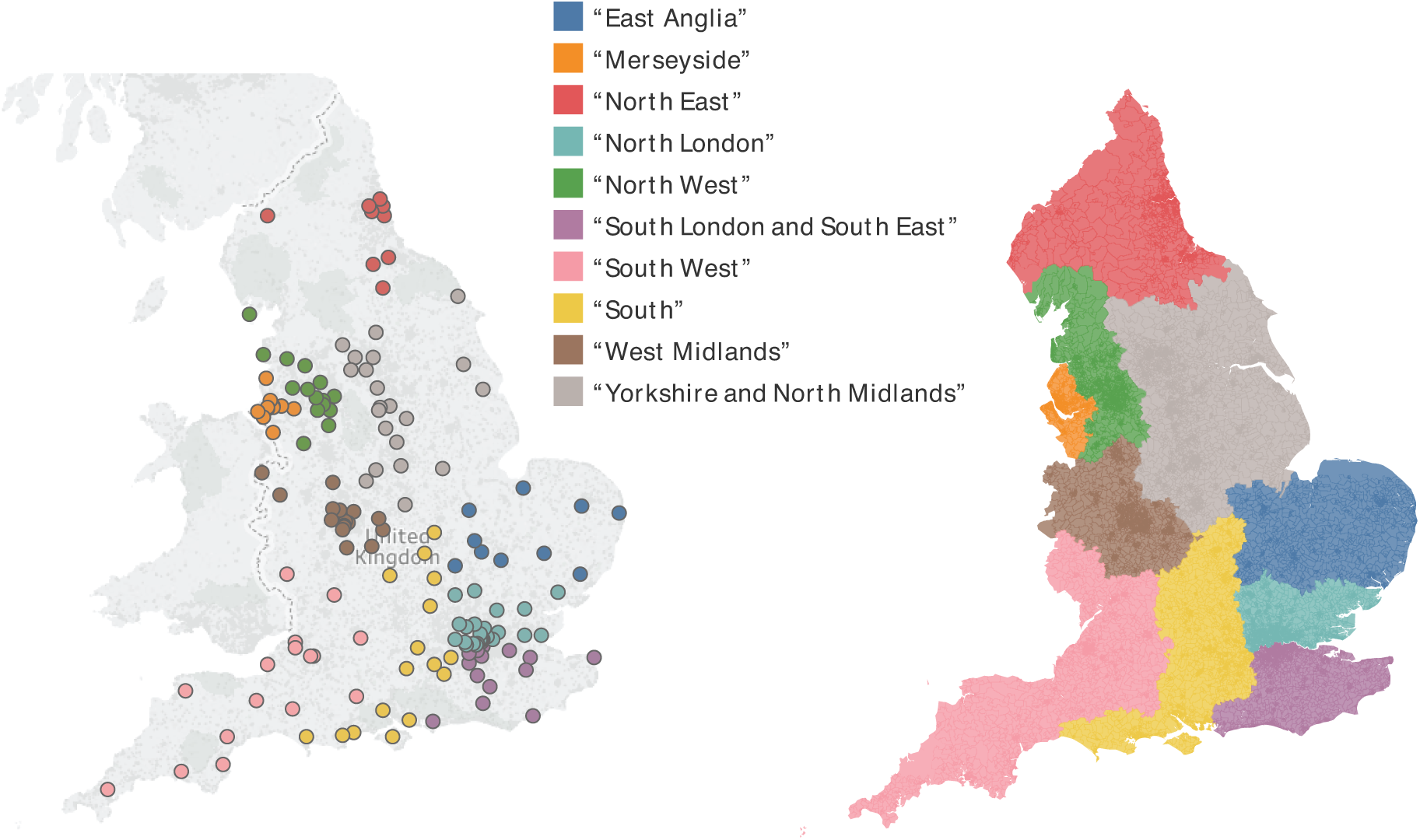
Spatial representation of data-sharing communities. The left-hand figure shows the geographic location of trusts coloured according to their assigned community. The right-hand figure shows the areas of the country where hospitals in each community are the most frequently attended. The 10 communities of hospitals obtained through community detection analysis of the patient-sharing network exhibit a strong geographical character.

## Discussion

To improve inter-organisational communication during transitions of care, it is critical to identify how the complete clinical record of a patient is distributed across care providers. This retrospective observational study used administrative data to identify the frequency and distribution of patient sharing between public hospital organisations in the NHS in England. Identification of hospital episodes involving 19.7 million patients over a 12-month period facilitated analysis of inter-organisational connections in England based on the patient-sharing network.

Of patients accessing secondary care on more than one occasion during the study period, 4.2 million patients (26.0%) presented to more than one provider. This high frequency of patient-sharing highlights the need for effective communication of patient information between healthcare organisations. On 14.7 million occasions (11.7%), patients who presented more than once to hospital had previously attended a different provider on their last hospital interaction, equivalent to 40,200 times per day, every day across England. Information pertaining to the care episode immediately prior to the next is likely to be of particular significance to patient management decisions. If this patient information is not available to providers due to poor inter-organisational communication, patients may be put at risk of transition of care errors ^7^.

Community detection methods identified groups of hospital organisations that regularly share patients and, therefore, would benefit most from improved data sharing with one another. Importantly, the process of identification of such patient-sharing communities of hospitals is data-driven and based on patterns of patient behaviour, rather than imposed by the presence of *a priori* administrative, geographic or historic relationships between hospitals. Unsurprisingly, many of the identified communities follow geographical regions and some will already have some degree of health information exchange. However, many existing regional inter-organisational data-sharing associations may be based on historical and service relationships, and the findings from this study provide empirical data on patient-sharing that may guide future health information exchange strategies. Improving interoperability within the ten communities identified may be particularly beneficial to enhancing health information exchange. Further research into the characteristics of patients being shared within these communities and the type of health information requiring exchange between them may provide further insights to guide specific service improvements.

The geographical mapping of inter-organisational patient sharing using Lower Layer Super Output Areas (LSOAs) illustrates the increased frequency of patient-sharing in areas with a higher spatial density of hospital trusts. Factors contributing to this may include centralisation of care between multiple institutions, ease of patient access to different closely located hospitals, or difficulty accessing care in previously attended hospitals due to bed shortages or other service issues.

Currently in England, there is significant variation in the type and use of health record systems between geographical regions and even between departments within hospitals^12^. Patients may have a mixture of electronic and paper records spread across several settings including primary care, secondary care, and radiology and pathology services. NHS England has aimed to connect electronic health records across settings by 2020^282930^. However, there are challenges associated with the implementation of shared electronic records^3132^ and establishing effective interoperability between existing health record systems^1133^. Poor interoperability is not unique to the UK, with a recent study in the United States finding that just 18.7% of hospitals report “often” using patient data from outside providers to inform patient care decisions^11^. The challenges of interoperability between EHR systems are well known to the NHS and the findings from this study should further encourage efforts to push for improvement. Approaches initially focused on developing local and regional patient information exchange within the patient-sharing communities identified in this study may improve the cost-effectiveness of health information exchange improvements.

This was a retrospective observational study using HES data from 2013 to 2015. Use of this dataset has facilitated a comprehensive overview of inter-organisational patient sharing in England. It is important to consider changes to organisational structures and systems in addition to evolving regional demographics in the period since collection of this data when applying these findings to the current population.

During the data collection period of this study, the organisation of health care in England underwent substantial change, in the transition from primary care trusts to clinical commissioning groups^34^. Regional boundaries for commissioning groups such as Local Area Teams and Specialised Commissioning Hubs established in 2012 took account of local geographies, service patterns and relationships at that time^35^. Geographic differences between those defined regional boundaries and the actual distribution of patient-sharing networks identified in this study are important to consider. Significantly, the community detection process used to identify the ten patient sharing communities shown in Figure 3 is based solely on patterns of patient sharing between providers obtained from patient-level administrative data, and therefore has no prior understanding of the organisational structure of secondary care. Where there are differences between the geographic distribution of service commissioning and the identified patient-sharing networks, this may indicate opportunities to modify the organisation of secondary care services to better reflect patterns of utilisation.

This paper has focused on inter-organisational patient-sharing connections. Previous, studies have identified significant heterogeneity within patient sharing networks with certain actors, whether hospitals, or individual physicians, exercising different roles within a network^222324^. Further analysis of the networks studied in this paper may offer additional insights into patient sharing within the NHS and further guide system improvements.

Patients in England frequently receive hospital care at two or more hospitals. Sharing of clinical information between providers is therefore an essential requirement of a safe, effective healthcare system. Through network analysis of hospital administrative data, this study has provided new insights into the distribution and frequency of patient sharing between hospital organisations in England. Through partitioning the national patient sharing network of England into 10 data-sharing communities, we have formulated a data-driven blueprint for the implementation of effective regional data-sharing communities in the English National Health Service. These methods may inform local, regional and national interoperability efforts in healthcare settings across the world with the aim of regionally interoperable, patient centred data sharing to drive safe, effective hospital care.

## Methods

### Data Processing and Data Sources

We performed a retrospective analysis of all patients receiving hospital care using non-public data from Hospital Episode Statistics provided by NHS Digital in England. Patients were identified from patient-level outpatient, inpatient and accident and emergency records from April 2013 to March 2015. Data were provided in a patient-level, pseudo-anonymised form. Interactions with hospital care are defined as a recorded presentation by a patient to emergency care, inpatient care or outpatient care.

All adult patients resident in England with a recorded interaction with secondary care in England in the year from 1st April 2013 to 31st March 2014 were included in the study. HES records were examined between 1^st^ April 2013 and 31^st^ March 2015 for subsequent interactions of these patients with secondary care in the 12 months following their first recorded presentation within the study period. A 12-month time window in which to identify instances of patient sharing was imposed to restrict instances of patient sharing to a clinically relevant interval of one year or less.

Providers were identified by the trust-level 3-digit provider code (PROCODE3), and therefore reflect hospital trusts rather than individual sites. It is thus possible for multiple hospital sites within the same trust to be represented by a single provider code. Where trusts cover multiple sites, the largest site according to number of inpatient beds was used as the location of the trust. To accommodate organisational change, providers that merged or separated over the period of study were treated as a single provider across the whole period.

### Descriptive Analysis

All patients presenting more than once to hospital within one year of their first recorded presentation in the study period were identified. Each interaction with hospital care for these patients was examined to determine the frequency with which two consecutive attendances involved different care providers, referred to as *fragmented presentations*. For each hospital, we calculated the proportion of clinical encounters involving fragmented presentations.

The Lower Layer Super Output Area (LSOA), a geographic division of England into 32844 distinct areas with an average population of 1614 people, analogous to United States Census block groups was used to map the patient encounters ^36^. For each LSOA, we calculated the median proportion of presentations that were fragmented. These values were mapped to generate an understanding of the relative frequency of patient sharing at a national level.

### Network Analysis

For each care provider, we calculated the number of presentations when a patient’s previous clinical encounter had been with each of the other providers. For each pair of care providers the number of occasions where consecutive presentations by a patient featured both providers, in either order, was calculated (*F*_*AB*_). Therefore, for every pair of providers in the study, we obtain a quantitative measure of the number of occasions where the most up-to-date record of a patient’s clinical care is held at one provider when they attend the other. These measures are then converted into a weighted, undirected network consisting of healthcare providers as nodes connected to one another by edges with weights *F*_*AB*_, and is termed the *patient-sharing network*.

Community detection based on modularity optimisation^37,38^ was then applied to identify stable communities of providers in an *unsupervised manner*, solely based on their weighted connectedness within the patient-sharing network. The communities obtained revealed a strong geographic linkage, so that we assigned a descriptive name to each community *a posteriori* (indicated with inverted commas) according to the approximate geographic location of the providers contained within it. The proportion of previous presentations to providers within the communities was compared to presentations to providers outside of the communities.

Figures 1 and 3 are created using Tableau version 2018.1 (Tableau Software, Seattle, USA) and figure 2 is created using Gephi version 0.9.2 (www.gephi.org). Map data copyrighted OpenStreetMap contributors and available from www.openstreetmap.org. This study received local ethical approval through the Imperial College Research Ethics Committee (17IC4178). As this was a national, retrospective study of routinely collected administrative data, informed consent from each human participant was not required.

### Data Availability

The hospital and LSOA constituents of each data-sharing community are available from the authors on request.

### Code Availability

Code written in Python version 3.6 is available upon request from the authors. Code is written specifically to analyse Hospital Episode Statistics in England.

## Acknowledgements

This article is independent research supported by grants from The Peter Sowerby Foundation and the National Institute for Health Research (NIHR) Imperial Patient Safety and Translational Research Centre (PSTRC). Infrastructure support for this work was provided by the NIHR Imperial Biomedical Research Centre (BRC). MB acknowledges support from EPSRC grant EP/N014529/1 supporting the EPSRC Centre for Mathematics of Precision Healthcare. The views expressed in this publication are those of the author(s) and not necessarily those of the NHS, the National Institute for Health Research or the Department of Health. The authors declare no competing interests in regard to this study.

## Author contributions

JC and LW were involved in all aspects of the study. SA, MB and AD were involved in the planning, interpretation, writing and reviewing of the study.

